# Genome-wide association study identifies 48 common genetic variants associated with handedness

**DOI:** 10.1101/831321

**Authors:** Gabriel Cuellar Partida, Joyce Y Tung, Nicholas Eriksson, Eva Albrecht, Fazil Aliev, Ole A Andreassen, Inês Barroso, Jacques S Beckmann, Marco P Boks, Dorret I Boomsma, Heather A Boyd, Monique MB Breteler, Harry Campbell, Daniel I Chasman, Lynn F Cherkas, Gail Davies, Eco JC de Geus, Ian J Deary, Panos Deloukas, Danielle M Dick, David L Duffy, Johan G Eriksson, Tõnu Esko, Bjarke Feenstra, Frank Geller, Christian Gieger, Ina Giegling, Scott D Gordon, Jiali Han, Thomas F Hansen, Annette M Hartmann, Caroline Hayward, Kauko Heikkilä, Andrew A Hicks, Joel N Hirschhorn, Jouke-Jan Hottenga, Jennifer E Huffman, Liang-Dar Hwang, Mohammad A Ikram, Jaakko Kaprio, John P Kemp, Kay-Tee Khaw, Norman Klopp, Bettina Konte, Zoltan Kutalik, Jari Lahti, Xin Li, Ruth JF Loos, Michelle Luciano, Sigurdur H Magnusson, Massimo Mangino, Pedro Marques-Vidal, Nicholas G Martin, Wendy L McArdle, Mark I McCarthy, Carolina Medina-Gomez, Mads Melbye, Scott A Melville, Andres Metspalu, Lili Milani, Vincent Mooser, Mari Nelis, Dale R Nyholt, Kevin S O’Connell, Roel A Ophoff, Cameron Palmer, Aarno Palotie, Teemu Palviainen, Guillaume Pare, Lavinia Paternoster, Leena Peltonen, Brenda WJH Penninx, Ozren Polasek, Peter P Pramstaller, Inga Prokopenko, Katri Raikkonen, Samuli Ripatti, Fernando Rivadeneira, Igor Rudan, Dan Rujescu, Johannes H Smit, George Davey Smith, Jordan W Smoller, Nicole Soranzo, Tim D Spector, Beate St Pourcain, John M Starr, Hreinn Stefánsson, Stacy Steinberg, Maris Teder-Laving, Gudmar Thorleifsson, Kari Stefansson, Nicholas J Timpson, André G Uitterlinden, Cornelia M van Duijn, Frank JA van Rooij, Jaqueline M Vink, Peter Vollenweider, Eero Vuoksimaa, Gérard Waeber, Nicholas J Wareham, Nicole Warrington, Dawn Waterworth, Thomas Werge, H.-Erich Wichmann, Elisabeth Widen, Gonneke Willemsen, Alan F Wright, Margaret J Wright, Mousheng Xu, Jing Hua Zhao, Peter Kraft, David A Hinds, Cecilia M Lindgren, Reedik Magi, Benjamin M Neale, David M Evans, Sarah E Medland

## Abstract

Handedness, a consistent asymmetry in skill or use of the hands, has been studied extensively because of its relationship with language and the over-representation of left-handers in some neurodevelopmental disorders. Using data from the UK Biobank, 23andMe and 32 studies from the International Handedness Consortium, we conducted the world’s largest genome-wide association study of handedness (1,534,836 right-handed, 194,198 (11.0%) left-handed and 37,637 (2.1%) ambidextrous individuals). We found 41 genetic loci associated with left-handedness and seven associated with ambidexterity at genome-wide levels of significance (P < 5×10^−8^). Tissue enrichment analysis implicated the central nervous system and brain tissues including the hippocampus and cerebrum in the etiology of left-handedness. Pathways including regulation of microtubules, neurogenesis, axonogenesis and hippocampus morphology were also highlighted. We found suggestive positive genetic correlations between being left-handed and some neuropsychiatric traits including schizophrenia and bipolar disorder. SNP heritability analyses indicated that additive genetic effects of genotyped variants explained 5.9% (95% CI = 5.8% – 6.0%) of the underlying liability of being left-handed, while the narrow sense heritability was estimated at 12% (95% CI = 7.2% – 17.7%). Further, we show that genetic correlation between left-handedness and ambidexterity is low (r_g_ = 0.26; 95% CI = 0.08 – 0.43) implying that these traits are largely influenced by different genetic mechanisms. In conclusion, our findings suggest that handedness, like many other complex traits is highly polygenic, and that the genetic variants that predispose to left-handedness may underlie part of the association with some psychiatric disorders that has been observed in multiple observational studies.

## Introduction

Handedness is the preferential use of one hand over the other. Hand preference is first observed during gestation as embryos begin to exhibit single arm movements [1, 2]. Across the life-span the consistent use of one hand leads to alterations in the macro- and micro-morphology of bone [3] resulting in enduring asymmetries in bone form and density [4, 5]. At the neurological level, handedness is associated with lateralization of language (the side of the brain involved in language) and other cognitive effects [6, 7]. The prevalence of left-handedness in modern western cultures is approximately 9% [8] and is greater in males than females [9]. While handedness is conceptually simple, its aetiology and whether it is related to brain and visceral (internal organs) asymmetry is unclear.

Since the mid-1980s, the literature regarding the genetics of handedness and lateralization has been dominated by the Right-shift [10] and Dextral-chance [11] theories. Both theories involve additive biallelic monogenic systems in which an allele at the locus biases an individual towards right handedness, while the second allele is a null allele that results in random determination of handedness by fluctuating asymmetry. The allele frequency of the right-shift variant has been estimated at ~43.5% [13], whilst that of the Dextral-chance variant has been estimated at ~20% in populations with a 10% prevalence of left-handedness[14]. A joint-analysis of data from 35 twin studies found that additive genetic factors accounted for 25.5% (95% CI: 15.7, 29.5%) of the phenotypic variance of handedness [12], which is consistent with predictions of the variance explained under the single gene right-shift and Dextral-change models. However, linkage studies [13–16] and genome-wide association studies (GWAS) [17–21] have failed to identify any putative major gene for handedness.

Most recently, two large-scale GWAS identified four genomic loci containing common variants of small effect associated with handedness [20, 21]. However, both GWAS failed to replicate signals at the *LRRTM1*, *PCSK6* and the X-linked Androgen receptor genes that had been reported previously in smaller genetic association studies [17–19]. In this study, we present findings from the world’s largest GWAS meta-analysis of handedness to date (N = 1,766,671), combining data from 32 cohorts from the International Handedness Consortium (IHC) (N = 125,612), 23andMe (N = 1,178,877) and UK Biobank (N = 462,182).

## Results

### Genome-wide association study of left-handedness

Across all studies, the handedness phenotype was assessed by questionnaire that evaluated either which *hand was used for writing* or for *self-declared handedness*. All cohorts were randomly ascertained with respect to handedness. Combining data across the 32 IHC cohorts, 23andMe and UK Biobank yielded 1,534,836 right-handed and 194,198 left-handed (11.0%) individuals [Supplementary Table 1]. After quality control (Methods), the GWAS meta-analysis included 13,346,399 SNPs (including autosomal and X chromosome SNPs) with a minor allele frequency (MAF)>0.05%.

The genetic correlations as estimated by bivariate LD-score regression [22] between the results from the UK Biobank, 23andMe and IHC GWAS were (r_g_^UKB-23andMe^ = 0.88, s.e. = 0.05, r_g_^UKB-IHC^ = 0.73, s.e. = 0.16, r_g_^IHC-23andMe^ = 0.60, s.e. = 0.11) suggesting that the three GWAS were capturing many of the same genetic loci for handedness. There was some inflation of the test statistics following meta-analysis (*λ*_GC_ = 1.22); however the intercept from LD-score regression analysis [23] was 1.01 suggesting that the inflation was due to polygenicity rather than bias due to population stratification or duplication of participants across the UK Biobank, 23andMe and IHC studies.

We identified 41 loci that met the threshold for genome-wide significance (P < 5×10^−8^) (Figure 1, Supplementary Table 2). Loci were defined as distinct if independent genome-wide significant signals were separated by at least 1Mb except for the MHC region and the 17q21.31 (that contains a common inversion polymorphism [24]) where we only report the lead signals due to the extent of linkage-disequilibrium in these loci. Summary statistics for the lead variants at genome-wide significant loci are presented in Table 1 along with the gene nearest to the lead SNP. A description of the putative functions of the nearest gene is included in Supplementary Table 2. Conditional analyses identified 9 additional independent SNPs at genome-wide significance near the lead SNPs on chromosome 2q, 6p, 16q and 17q [Supplementary Table 3]. Interestingly, the list of genome-wide significant associations included multiple variants close to genes involved in microtubule formation or regulation (*i.e. MAP2, TUBB, TUBB3, NDRG1, TUBB4A, TUBA1B, BUB3, TTC28*). A phenome-wide association scan (PheWAS) of the lead SNPs using GWAS summary data from 1,349 traits revealed that 28 out of the 41 lead SNPs have been previously associated with other complex traits (Supplementary Table 4). Among these results, we highlight that the rs6224, rs13107325 and rs45527431 variants have been previously associated with schizophrenia at genome-wide levels of significance (P < 5×10^−8^). Specifically, the effect of alleles associated to both traits had the same direction of effect (e.g. those that increase odds of left-handedness also increase risk of schizophrenia) for all three SNPs. Further, we found that seven variants associated with left-handedness were also associated with educational attainment; however, the direction of effects of these SNPs on left-handedness and educational attainment was not consistent. Future colocalization will be needed to assess if the same SNP associated to handedness affects these traits of is a product of correlation between causal SNP.

**Table 1.**
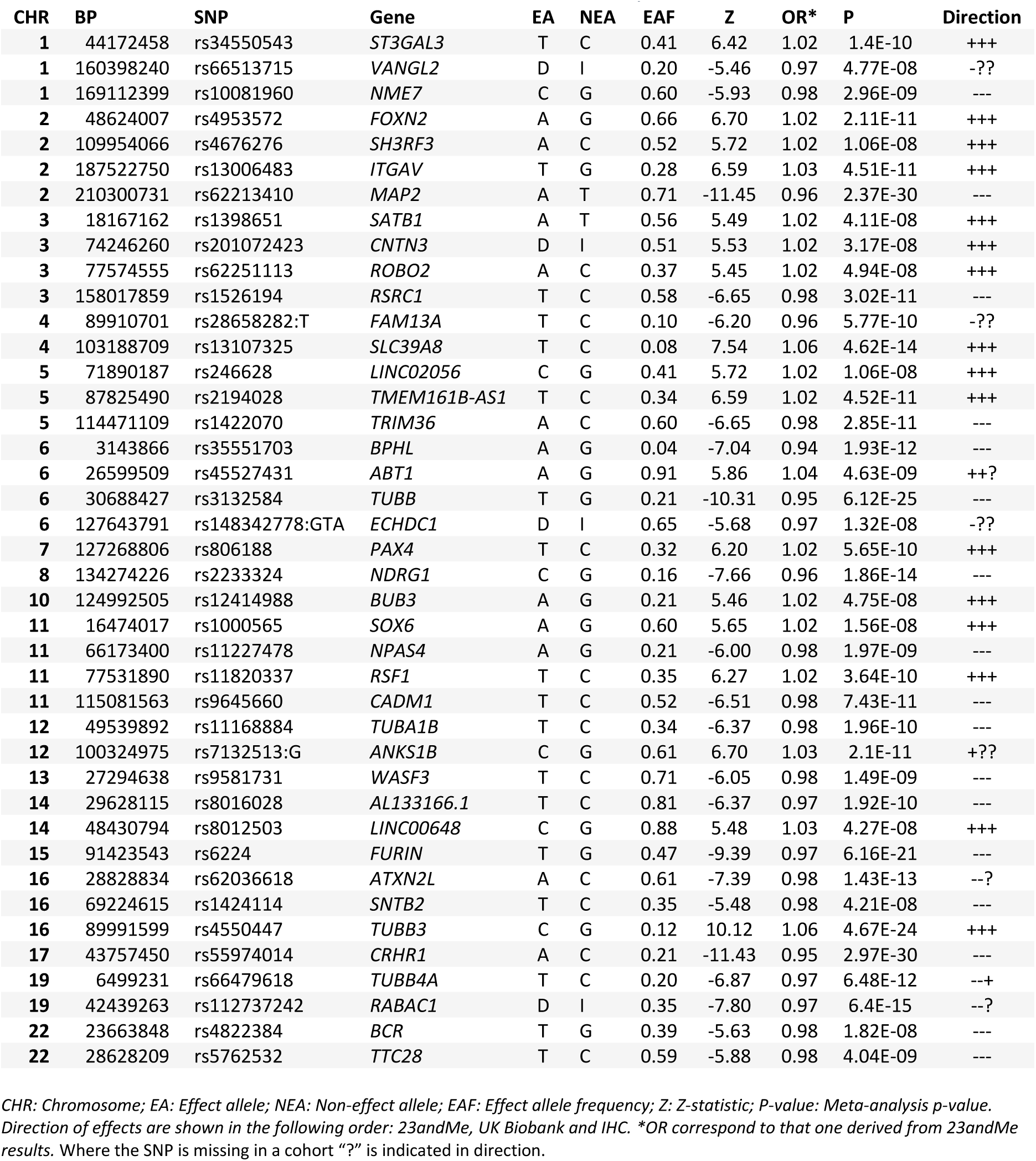
Loci associated with left-handedness after meta-analysis of 23andMe, UK Biobank and IHC.

**Figure 1.**
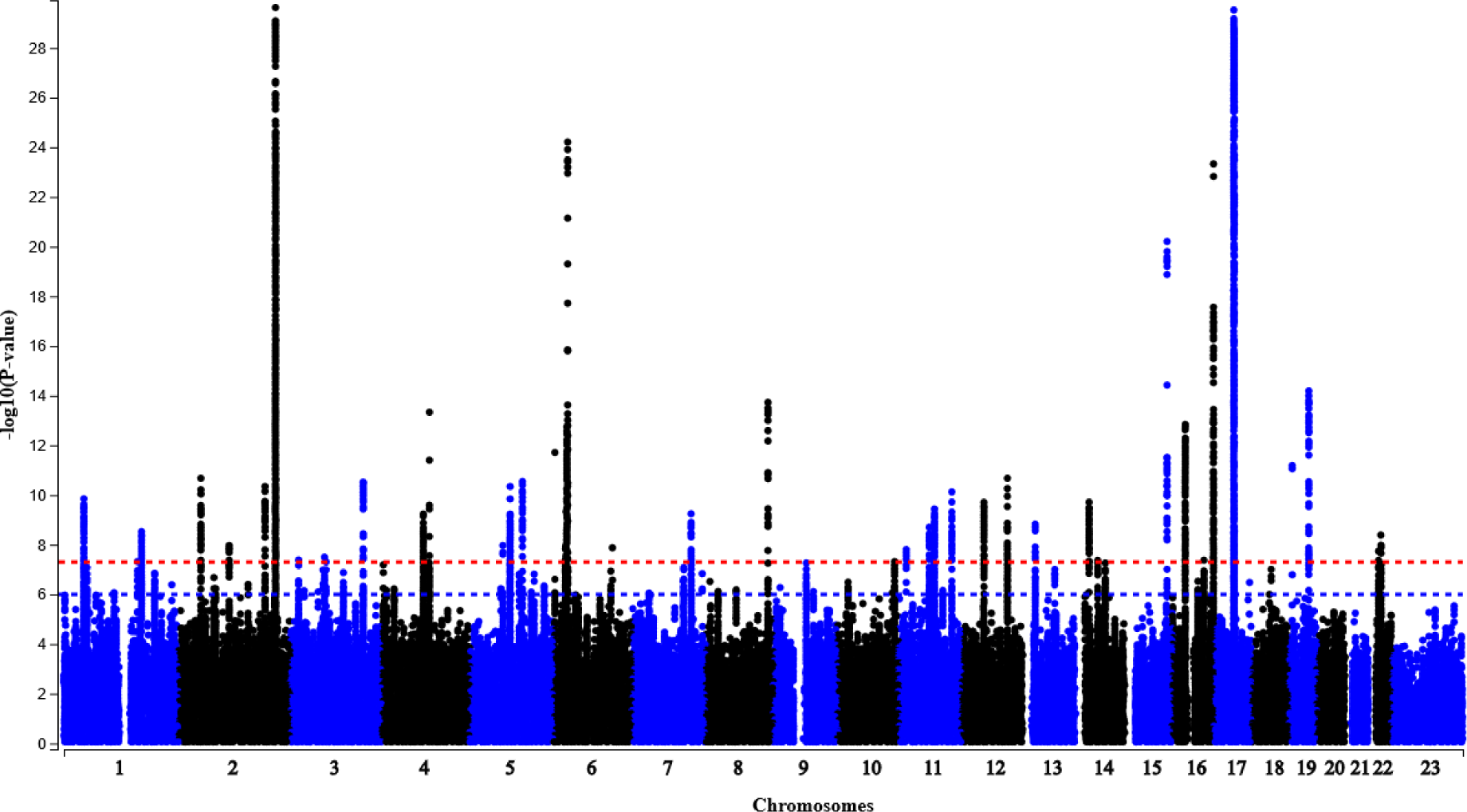
Manhattan plot of left-handedness Meta-analysis

In order to identify the most likely tissues and pathways underling the association of the SNPs, we used DEPICT [25]. Results from tissue enrichment analysis implicated the central nervous system, including brain tissues such as the hippocampus and cerebrum at FDR < 5% (Table 2), consistent with the hypothesis that handedness is primarily a neurological trait. We observed statistically significant pathways (FDR < 5%) including microtubule regulation and cerebral cortex and hippocampus morphology (Table 3). Inter alia, we then performed gene-based analyses using gene-expression prediction models of brain tissues using S-MultiXcan to identify additional loci [26]. In total, we tested the association between the predicted expression of 14,501 genes in brain tissues and left-handedness. In addition to detecting significant associations (P < 3.44×10^−6^) of genes within the loci identified during the meta-analysis, we observed an association between left-handedness and the predicted expression of *AMIGO1* (P = 2.82 × 10^−7^), a gene involved in growth and fasciculation of neurites from cultured hippocampal neurons and that may be involved in myelination of developing neural axons [27]. Supplementary Table 5 shows all the statistically significant associations from the S-MultiXcan analysis.

**Table 2.** Results of the tissue enrichment analysis for left-handedness (DEPICT). Only results with FDR < 5% are shown.

| **Tissue** | **Group** | **P-value** |
| --- | --- | --- |
| **Corpus Striatum** | Nervous System | 0.00000673 |
| **Basal Ganglia** | Nervous System | 0.0000148 |
| **Hippocampus** | Nervous System | 0.0000227 |
| **Central Nervous System** | Nervous System | 0.0000327 |
| **Brain** | Nervous System | 0.0000357 |
| **Telencephalon** | Nervous System | 0.0000398 |
| **Parahippocampal Gyrus** | Nervous System | 0.0000414 |
| **Entorhinal Cortex** | Nervous System | 0.0000414 |
| **Limbic System** | Nervous System | 0.0000423 |
| **Cerebrum** | Nervous System | 0.0000427 |
| **Prosencephalon** | Nervous System | 0.0000487 |
| **Temporal Lobe** | Nervous System | 0.0000587 |
| **Cerebral Cortex** | Nervous System | 0.0000668 |
| **Parietal Lobe** | Nervous System | 0.000305 |
| **Mesencephalon** | Nervous System | 0.000843 |
| **Occipital Lobe** | Nervous System | 0.00131 |
| **Visual Cortex** | Nervous System | 0.00156 |
| **Brain Stem** | Nervous System | 0.00166 |

**Table 3.**
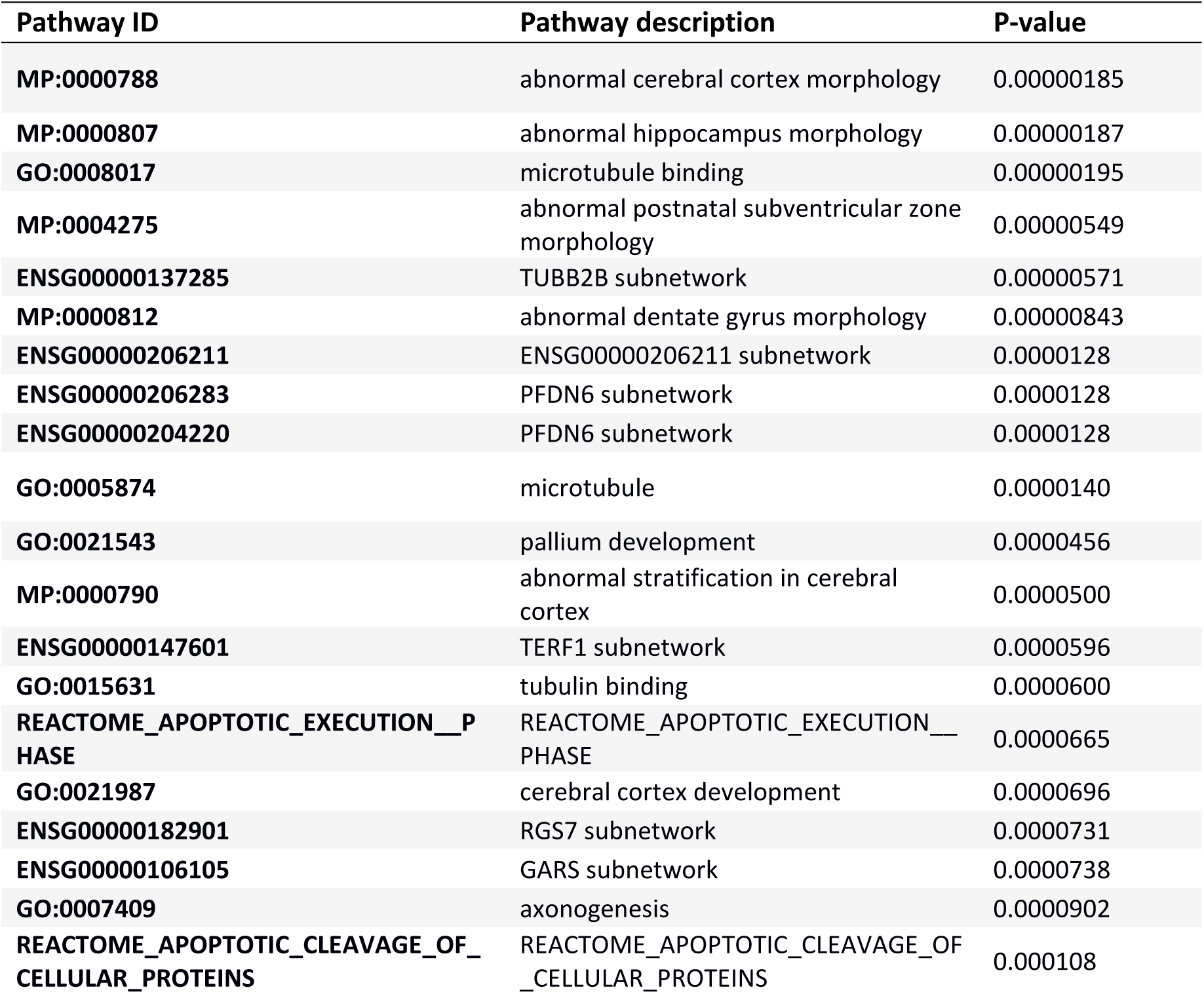
Results from pathway enrichment analysis for left-handedness (DEPICT). Only results with FDR < 5% are shown.

Multiple studies have reported that left-handedness and ambidexterity are more prevalent in males than in females [9]. Consistent with this observation, we found that 11.9% of male participants in the IHC cohorts reported being left-handed or ambidextrous, compared to only 9.3% of females (OR = 1.31; 95% CI: 1.25 – 1.38; P < 2.2×10^−16^) [Supplementary Table 6]. Similarly, in UK Biobank, 10.5% of males and 9.9% of females were left handed (OR = 1.07; 95% CI: 1.05 – 1.09; P = 1.87×10^−11^) and in 23andMe 15.6% of males and 12.6% of females were left handed (OR = 1.28; 95% CI: 1.26 – 1.30; P < 2.2×10^−16^). Sex differences in ambidexterity were also apparent in the UK Biobank and 23andMe cohorts. In UK Biobank 2% of males and 1.30% of females reported being ambidextrous (OR = 1.55; 95% CI: 1.47 – 1.62; P < 2.2×10^−16^) and in 23andMe 3.45% of males and 2.61% of females (OR = 1.33; 95% CI: 1.28 – 1.37; P < 2.2×10^−16^). Birth year had a small but significant effect on left-handedness with individuals who were born more recently being more likely to be left-handed (OR = 1.008 per year; 95% CI: 1.007 – 1.009; P < 2.2×10^−16^).

The differences in prevalence between males and females and a previously reported association between the X-linked androgen receptor gene and handedness [22] could reflect the involvement of hormone-related genes in handedness aetiology. We therefore carried out a sex-stratified GWAS of handedness in the UK Biobank data using left-handed individuals as cases and right-handed individuals as controls; however, we did not identify any genome-wide significant loci. Despite this, the point estimate of the genetic correlation between male handedness and female handedness computed using LD-score regression was lower than unity but not significantly different from one (r_G_ = 0.77, s.e. = 0.12, P = 0.055).

### Associations with previously reported candidate genes

All loci identified in recent GWAS of handedness from UK Biobank, replicated [20, 21]. However, we found no evidence of association between left-handedness and genes and genetic variants reported in prior studies. The SNPs rs1446109, rs1007371 and rs723524 in the *LRRTM1* locus reported by Francks *et al* [18] did not reach nominal significance in any of the analyses performed (P>0.05). Similarly the SNP rs11855415 reported by Scerri *et al* [19] as associated with left-handedness in dyslexic individuals also did not exhibit an association (P>0.05). Further, we investigated if the 27 genes exhibiting asymmetric expression in early development of the cerebral cortex described by Sun *et al* [28] were also associated with handedness in our S-MultiXcan analyses. Only 11 out of the 27 asymmetry genes were available in our analysis, and after adjusting the results for multiple testing, we did not observe any significant association [Supplementary Table 7]. In a more recent study, Ocklenburg and colleagues [29] list 74 genes displaying asymmetric expression in cervical and anterior thoracic spinal cord segments of five human foetuses. In total, 43 out of the 74 genes were in our S-MultiXcan analyses, of which only *HIST1H4C* was statistically significant after correcting for multiple testing (P = 2.2×10^−4^) [Supplementary Table 8].

### Heritability of left-handedness and genetic correlations with other traits

Twin studies estimate the heritability of left-handedness as around 25% [12]. In the present study, we employed a diverse range of statistical methods including LD-score regression, G-REML analysis as implemented in BOLT-LMM, and maximum likelihood analysis of identity by descent sharing in close relatives [30] to provide complementary estimates of SNP heritability and total heritability that relied on a different set of assumptions than the classical twin model. Using GWAS summary statistics from our study and LD-score regression, we estimated that the variance explained by SNPs was 3.45% (s.e. = 0.17%) on the liability scale with prevalence of left-handedness set at 10% [Table 4]. Using genotypic data from the UK Biobank Study (and age and sex as covariates) and G-REML analysis, we also obtained low estimates of the SNP heritability (5.87%, s.e. = 2.21%). Due to the large disparity between estimates of heritability from twin studies and the lower estimates of SNP heritability from the above approaches, we estimated the heritability of handedness using the autosomal Identity By Descent (IBD) information from closely related individuals [28] in the UK Biobank (estimated genome-wide IBD>8%). We partitioned the phenotypic variance into additive genetic effects (A), shared environmental effects (C) and individual environmental inputs (E) (see Methods section). We estimated that additive genetic effects explained 11.9% (95% CI: 7.2 – 17.7) of the phenotypic variance in handedness while shared environment and individual environment accounted for 4.6% (95% CI: 0 – 9.0) and 83.6% (95% CI: 75.2 – 85.6) of the variance in liability respectively [Table 4]. Dropping (C) from the model did not significantly worsen the fit of the model (P=0.29). Estimates from the A+E model were 19.7% (95% CI: 13.6 – 25.7) for additive genetic effects, overlapping with those from twin studies.

**Table 4.** SNP heritability and heritability of left-handedness estimated using a range of different approaches.

| *Data Used* | *Method* | *$h_g^2$ (s.e.) – liability scale* |
| --- | --- | --- |
| *IHC Meta-analysis (32 studies)* | LD-score regression | 0.031<br>(0.013) |
| *UK Biobank left-handed individuals only as cases* | LD-score regression | 0.033<br>(0.004) |
| *23andMe left-handed individuals only as cases* | LD-score regression | 0.040<br>(0.002) |
| *Meta-analysis UK biobank, 23andMe and IHC* | LD-score regression | 0.035<br>(0.002) |
| *UK Biobank left-handed individuals only as cases (males)* | LD-score regression | 0.042<br>(0.006) |
| *UK Biobank left-handed individuals only as cases (females)* | LD-score regression | 0.032<br>(0.005) |
| *UK Biobank left-handed individuals only as cases* | REML (BOLT-LMM) | 0.059<br>(0.003) |
| *Right vs Left handed 0.08 < IBD < 0.3 Relatives (no C) + siblings 0.65 > IBD > 0.35 with C in the model.* | ACE model | A = 0.12*<br>(95% CI: 0.07 – 0.17)<br>C = 0.045<br>(95% CI: 0 – 0.09) |
| *Right vs Left handed 0.08 < IBD < 0.3 Relatives (no C) + siblings 0.65 > IBD > 0.35 without C in the model.* | AE model | A = 0.20<br>(95% CI: 0.14 – 0.26) |
| *Meta-analysis of twin studies of handedness [12]* | ACE model | A = 0.25<br>(95% CI: 0.157 – 0.30)<br>C = 0 (95% CI = 0 – 0.076%) |

\*Estimate of narrow sense heritability ( $h^2$ )

We investigated the genetic correlation between left-handedness and 1349 complex traits using LD-score regression as implemented in CTG-VL [31]. We did not observe any statistically significant genetic correlation at a Bonferroni P-value threshold (0.05/1349 = 3.7 × 10^−5^). We observed suggestive (P < 0.05) positive correlations with neurological and psychiatric traits including schizophrenia, bipolar disorder, intra-cranial volume and educational attainment and negative correlations with mean pallidum volume [Supplementary Table 9]. Interestingly, the genetic correlation between our left-handedness meta-analysis and our ambidexterity meta-analysis was only moderate (r_g_ = 0.24, s.e = 0.03), suggesting divergent genetic etiologies.

### Genome-wide association study of ambidexterity

Given the moderate genetic correlation between left-handedness and ambidexterity, a separate GWAS of ambidexterity was carried out using UK Biobank and 23andMe data using ambidextrous individuals as cases (N= 37,637; ~2% of the total sample) and right-handed individuals as controls (N=1,422,823). This meta-analysis included 12,493,443 autosomal and X chromosome SNPs with a MAF>0.05%.

Similar to the left-handedness GWAS, prior to meta-analysis, we computed the genetic correlation between the UK Biobank ambidexterity GWAS and the 23andMe GWAS. The estimate of the genetic correlation was r_g_ = 1 (s.e. = 0.15) indicating that both GWAS were capturing the same genetic loci. After meta-analysis, we identified seven loci with P < 5×10^−8^ (Figure 2). Table 5 displays the summary statistics for the lead SNPs at these loci along with the closest gene. Full summary statistics and description of the nearest gene for these loci are included in Supplementary Table 10. There was some overlap between genome-wide significant SNPs associated with left-handedness and ambidexterity with 16 out of the 41 SNPs associated to left-handedness displaying a nominal significant association with ambidexterity (P < 0.05); 15 of which were also in the same direction of effect (Supplementary Table 11). Conditional analyses did not identify further independent signals at genome-wide levels of significance. PheWAS revealed that the lead SNPs have been implicated in anthropometric traits and blood biomarkers [Supplementary Table 12].

**Table 5.** Loci associated with ambidexterity through meta-analysis of 23andMe and UK Biobank.

| CHR | BP | SNP | Gene | EA | NEA | EAF | Z | OR* | P | Direction |
| --- | --- | --- | --- | --- | --- | --- | --- | --- | --- | --- |
| 1 | 150317558 | rs782122127 | PRPF3 | D | I | 0.19 | -6.32 | 0.88 | 2.70E-10 | ?- |
| 2 | 58196110 | rs2030237 | VRK2 | A | G | 0.58 | 5.84 | 1.04 | 5.29E-09 | ++ |
| 2 | 104437850 | rs139630683 | AC013727.1 | D | I | 0.45 | 5.85 | 1.05 | 4.88E-09 | +? |
| 7 | 91899117 | rs2040498 | ANKIB1 | A | T | 0.65 | 6.21 | 1.06 | 5.42E-10 | ++ |
| 8 | 77104817 | rs10113066 | RNU2-54P | T | G | 0.51 | 5.90 | 1.05 | 3.74E-09 | ++ |
| 10 | 89722731 | rs36062478 | PTEN | T | C | 0.87 | -5.62 | 0.94 | 1.87E-08 | -- |
| 12 | 49530132 | rs35554786 | TUBA1B | D | I | 0.24 | -6.05 | 0.93 | 1.49E-09 | -? |

CHR: Chromosome; EA: Effect allele; NEA: Non-effect allele; EAF: Effect allele frequency; Z: Z-statistic; P-value: Meta-analysis p-value.
Direction of effects are shown in the following order: 23andMe and UK Biobank. \*OR correspond to that one derived from 23andMe results.
. Where the SNP is missing in a cohort “?” is indicated in direction.

**Figure 2.**
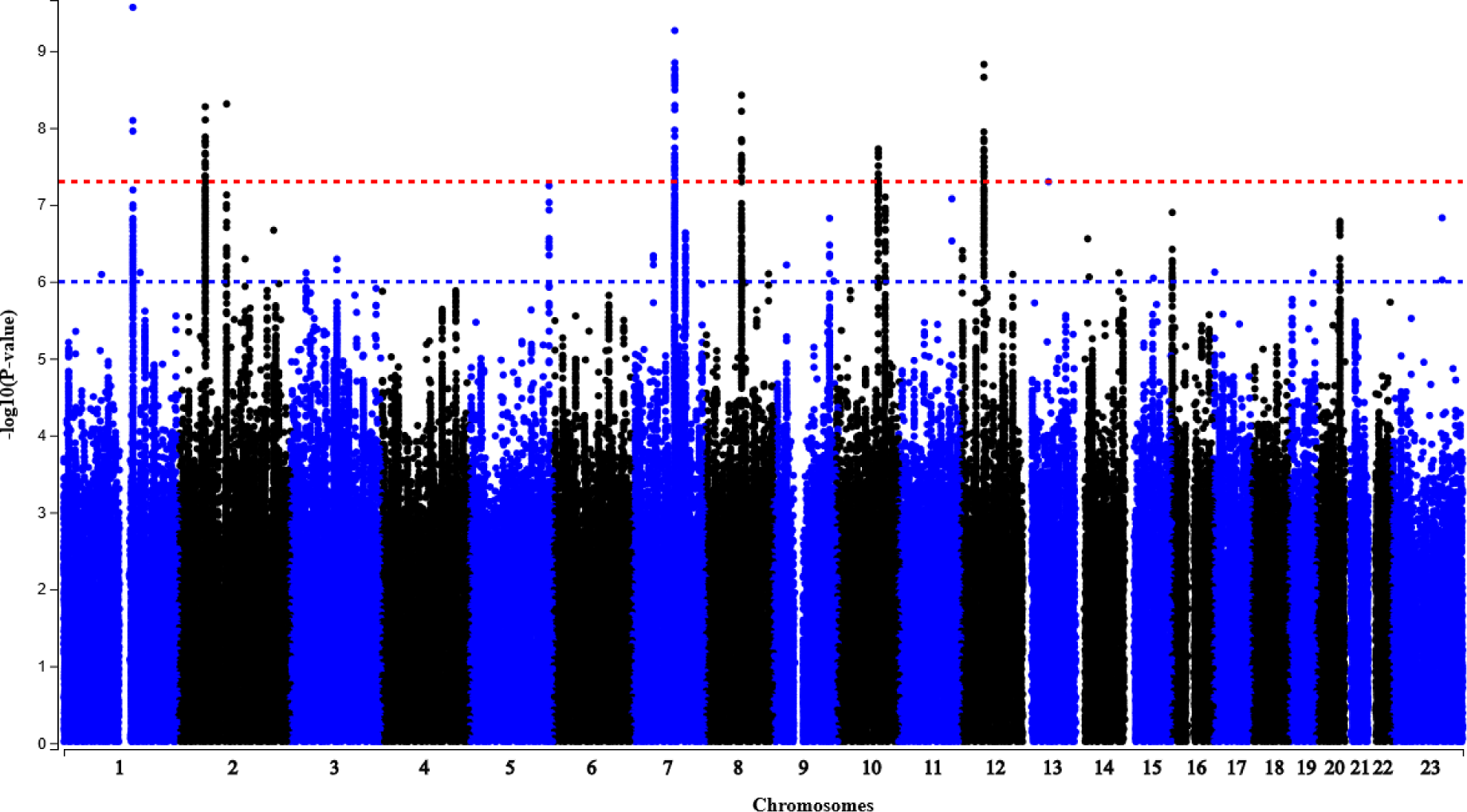
Manhattan plot of ambidexterity Meta-analysis

DEPICT analyses did not identify any tissue or pathway at an FDR < 5%. The top associated tissues were cartilage and smooth muscle (P = 0.016 and P = 0.025 respectively). Top tissue and pathway enrichment results are included in Supplementary Table 13 and 14. S-MultiXcan analyses based on the association between predicted gene-expression on brain tissues and ambidexterity identified the genes *QTRTD1*, *TMEM215*, *RPL41* and *RAB40C* in addition to those loci identified during the GWAS [Supplementary Table 15].

### Heritability of Ambidexterity and genetic correlations

Due to the low prevalence of ambidexterity in the population there are no heritability estimates available from twin studies. Similarly, the number of ambidextrous individuals in UK Biobank was not enough to precisely estimate the heritability using maximum likelihood analysis of identity by descent sharing in close relatives. However, the SNP heritability of ambidexterity on the liability scale (1% prevalence) estimated through LD-score regression and REML implemented in BOLT-LMM was higher than that observed for left-handedness (h^2^_g_ = 0.12 (s.e. = 0.007) and h^2^_g_ = 0.15 (s.e. = 0.014)).

We estimated the genetic correlation between ambidexterity and a catalogue of 1349 traits with GWAS summary statistics. Our analyses revealed 260 statistically significant (P < 3.7 × 10^−5^) genetic correlations. Among the strongest correlations, we saw positive genetic correlations between ambidexterity and traits related to pain and injuries, body mass index, as well as a negative genetic correlation with educational attainment [Supplementary Table 16].

## Discussion

We have carried out the largest genetic study of handedness to date. Our GWAS and SNP heritability analyses demonstrate conclusively that handedness is a polygenic trait – with multiple genetic variants increasing the odds of being left-handed or ambidextrous by a small amount. We identified 41 left-handedness and 7 ambidexterity loci that reached genome-wide significance. Our findings are in contrast to the single gene right-shift [13] and dextral-chance[14] hypotheses, where the causal genes are hypothesised to account for the heritability of handedness. If these large effect variants do exist, they should have been detected by our GWAS meta-analysis which provided over 90% statistical power [Supplementary Table 17] to detect variants with effect sizes as small as a 5% increase in odds per allele for common variants (MAF>0.05) at genome-wide significance (α = 5×10^−8^). Instead, the present findings firmly support the hypothesis that handedness, like many other behavioural and neurological traits, is influenced by many variants of small effect.

Using different methods and cohorts, we estimated the SNP heritability (h_g_^2^) of handedness to be between 3% and 6%. However, by using IBD-based methods applied to siblings and other relative pairs, we estimated the narrow sense heritability (h^2^) to be 11.9% (95% CI: 7.2 – 17.7). Although this is lower than that obtained from twin studies (25%; 95 CI: 15.7 – 29.5 [12] and 21%; 95% CI: 11 – 30 [32]), the confidence intervals for the estimates overlap. Interestingly, h_g_^2^ estimates for ambidexterity were larger (12%-15%), suggesting that common SNPs tag a higher proportion of variability in liability to ambidexterity than in liability to left-handedness.

Interestingly, our GWAS meta-analysis of left-handedness identified eight loci close to genes involved in microtubule formation and regulation. An enrichment for microtubule-related pathways was then confirmed by the DEPICT analysis. Microtubules are polymers that form part of the cytoskeleton and are essential in several cellular processes including intracellular transport, cytoplasmic organization and cell division. With respect to handedness, microtubule proteins play important roles during development and migration of neurons, plasticity, and neurodegenerative processes [33, 34]. The association between handedness and variation in microtubule genes also provides insights into differences in prevalence of various neuropsychiatric disorders and left-handedness observed in some epidemiological studies [35, 36]. Recent genetic studies have identified mutations in a wide variety of tubulin isotypes and microtubule-related proteins in many of the major neurodevelopmental and neurodegenerative diseases [33, 37–39].

We observed an association between left-handedness and the 17q21.31 locus. A deletion in this locus is known to cause Koolen de Vries syndrome, a disorder characterized by intellectual disability, developmental delay and neurological abnormalities of the corpus callosum, hippocampi and ventricles. Variation in the 17q21 locus, including structural variation, has been associated with schizophrenia [40], autism [41, 42] and cognition [43]. In addition, based on our pheWAS, the rs55974014 within this locus has been associated with mood swings, neuroticism and educational attainment traits. The rs55974014 SNP is located near several genes with neurological functions, including *CRHR1* and *NSF*. Other SNPs close to this gene have been associated with intelligence [44] and Parkinson’s disease [45]. Future colocalization analyses are warranted to assess the veracity of the same variant affecting multiple traits.

The genetic correlation between left-handedness and ambidexterity was low, suggesting that the genetic architecture underlying the two traits is different. Whereas being left-handed was not statistically significantly (genetically) correlated with any other trait among the 1,349 complex traits tested, ambidexterity was significantly correlated with multiple traits, particularly anthropometric and those involving pain and injuries. This suggests that reporting being able to write with both hands may be a result from injuries that led to use of the other hand. Future studies into the genetics of ambidexterity should include detailed phenotyping that considers the reasons leading to hand use preference. Among the suggestive genetic correlations (P < 0.05), we observed positive genetic correlations between left-handedness and schizophrenia and bipolar disorder consistent with some previous observations of greater atypical hand dominance in schizophrenia and bipolar disorder patients [46, 47].

The present study benefitted from having a large sample size that allowed the detection of dozens of novel variants of small effect on handedness. However, it is worth noting that the genetic correlations derived from GWAS summary statistics of left-handedness in the IHC, 23andMe and UK Biobank were high but statistically different from one, potentially impacting the statistical power of the meta-analysis. These differences may have been due to the way data was collected in each of the cohorts. For example, in the UK Biobank, handedness data was obtained at up to three occasions, while this was not the case for the 23andMe and the IHC cohorts. Furthermore, genetic correlations with IHC may have been affected by the IHC having combined ambidextrous and left-handed individuals as cases, and the fact that different imputation panels were used for the 32 cohorts that make up IHC.

In summary, we report the world’s largest GWAS meta-analysis of handedness. We showed that handedness is polygenic and found evidence that microtubule genes may play an essential role in lateralization. Loci mapped in the present study warrant further exploration of their potential role in neurological development and laterality.

## Methods

### Genome-wide association in UK Biobank

UK Biobank is a large long-term biobank study from the United Kingdom that aims to identify the contribution of genetic and environmental factors to disease. Detailed information on phenotyping and genotyping is presented elsewhere [48]. In brief, the UK Biobank recruited 502,647 individuals aged 37-76 across the country and gathered information regarding their health and lifestyle, including handedness via questionnaires. Genotype data from the UK Biobank is available for 487,411 participants. Genotypes were imputed by UK Biobank against the UK10K reference panel using IMPUTE 2 [49]. In addition to the quality control metrics performed centrally by UK Biobank [48], we defined a set of participants of European ancestry by clustering the first two principal components (PCs) derived from the genotype data. Using a K-means algorithm with K=4, we identified a group of 463,023 individuals of European ancestry. From this group, 462,182 individuals (250,767 females) had data on handedness. In total 410,677 participants identified themselves as right-handed, 43,859 as left-handed and 7,646 as ambidextrous. The mean birth year of the participants was 1951 (*s.d.* = 8.04).

We tested 11,498,822 autosomal and X chromosome SNPs with minor allele frequency (MAF>0.005) and info score >0.4 for association with handedness using BOLT-LMM which implements a linear mixed model to account for cryptic relatedness and population structure. Sex and age were included as covariates in all models. We performed four analyses 1) right- vs left-handed; 2) right vs ambidextrous; 3) right- vs left-handed (male only); and 4) right- vs left-handed (female only). Analyses of X chromosome genotypes were performed in BOLT-LMM fitting sex as a covariate and coding the male genotypes as 0/2.

### Genome-wide association in 23andMe

All individuals included in the analyses were research participants of the personal genetics company 23andMe, Inc., a private company, whose phenotypes were collected via online surveys. DNA extraction and genotyping were performed on saliva samples by the National Genetics Institute (NGI), a CLIA licensed clinical laboratory and a subsidiary of Laboratory Corporation of America. Samples were genotyped on one of five genotyping platforms. The v1 and v2 platforms were variants of the Illumina HumanHap550 + BeadChip, including about 25,000 custom SNPs selected by 23andMe, with a total of about 560,000 SNPs. The v3 platform was based on the Illumina OmniExpress+ BeadChip, with custom content to improve the overlap with the v2 array, with a total of about 950,000 SNPs. The v4 platform was a fully customized array, including a lower redundancy subset of v2 and v3 SNPs with additional coverage of lower-frequency coding variation, and about 570,000 SNPs. The v5 platform was an Illumina Infinium Global Screening Array (~640,000 SNPs) supplemented with ~50,000 SNPs of custom content. Samples that failed to reach 98.5% call rate were re-analysed. Individuals whose analyses failed repeatedly were re-contacted by 23andMe customer service to provide additional samples.

For our standard GWAS, we restricted participants to a set of individuals who have a specified ancestry (predominantly European ancestry) determined through an analysis of local ancestry [50]. A maximal set of unrelated individuals was chosen for each analysis using a segmental identity-by-descent (IBD) estimation algorithm [51]. Individuals were defined as related if they shared more than 700 cM IBD, including regions where the two individuals share either one or both genomic segments IBD. This level of relatedness (roughly 20% of the genome) corresponds approximately to the minimal expected sharing between first cousins in an outbred population.

We used Minimac3 to impute genotype data against a reference panel comprised of the May 2015 release of the 1000 Genomes Phase 3 haplotypes [52] and the UK10K imputation reference panel [53]. We computed associations by logistic regression assuming additive allelic effects. We used the imputed dosages rather than best-guess genotypes and included covariates for age, sex, the top five principal components to account for residual population structure, and indicators for genotype platforms to account for genotype batch effects. For associations on the X chromosome, male genotypes were coded as if they were homozygous diploid for the observed allele. For QC of the GWAS results, we removed SNPs with rsq < 0.3, MAF < 0.005 and available sample size <20% of the total sample, as well as SNPs that had strong evidence of a platform batch effect. We also flagged logistic regression results that did not converge due to complete separation, identified by abs(effect)>10 or s.e. >10 on the log odds scale.

### International handedness consortium

The International Handedness Consortium (IHC) is a large-scale collaboration between 32 cohorts (N=125,612) with existing genome-wide association (GWAS) data to identify common genetic variants influencing handedness. Across all studies, the phenotype was collected by questionnaire by asking either which *hand was used for writing* or for *self-declared handedness.* As these two measures are highly (~95%) concordant, and less than 1% of participants reported being able to write with both hands [8, 30] both left-handed and ambidextrous individuals were classified as cases. All cohorts were population samples with respect to handedness, thus combining the data from the 32 studies yielded 13,599 left-handed and 112,013 right-handed individuals [Supplementary Table 1].

All individuals were of self-declared European ancestry (confirmed by genotypic PCA in each cohort). Within each cohort, the genotypic data were imputed to Phase I and II combined HapMap CEU samples (build 36 release 22) with the exception of the Finnish Twin Cohort study, Health Professionals Follow-Up Study and Nurses’ Health Study HPFS/NHS, Netherlands Twin Registry (NTR), and TOP cohorts which were imputed to 1000G phase 3 V5 European population. Within each sample, genome-wide association analyses were conducted for both genotyped and imputed SNPs. The imputed genotypes were analysed using the dosage of an assumed effect allele under an additive model with covariates for year of birth and sex. Supplementary Table 18 show the imputation and analysis software used in each of the cohorts.

### Meta-analysis of IHC, UK Biobank and 23andMe

A weighted Z-score meta-analysis was conducted with the METAL software [54] using the summary GWAS statistics from each of the 32 IHC cohorts, UK Biobank and 23andMe. Given the large discrepancies between the number of cases and controls, we elected to weight each sample by the effective sample size for binary traits, defined as N_eff_=4/(1/N_cases_+1/N_controls_). Prior to meta-analysis, quality control thresholds were applied to each of the GWAS results from the individual studies (r^2^ ≥ 0.3, MAF ≥ 0.005, P_HWE_ ≥ 1×10^−5^). We also removed genetic variants for which the frequency substantially differed from one of the Haplotype Reference Consortium panels (frequency difference > 0.2). We used EasyQC [55] to identify SNPs that had allele frequencies which differed substantially from the Haplotype Reference Consortium. In total, up to 13,346,399 SNPs remained for the left-handedness meta-analysis. For the ambidexterity meta-analysis, only 23andMe and the UK Biobank were used. This meta-analysis included up to 12,493,443 SNPs.

### Tissue expression and pathway analyses

Tissue expression and pathway analyses were performed using DEPICT [25] implemented in the Complex-Traits Genetics Virtual Lab (CTG-VL) [31]. DEPICT assesses whether genes in associated loci are highly expressed in any of the 209 Medical Subject Heading (MeSH) tissue and cell type annotations based on RNA-seq data from the GTEx project [56]. Molecular pathways were constructed based on 14,461 gene sets from diverse database and data types, including Gene ontology, Kyoto encyclopedia of genes and genomes (KEGG) and REACTOME. Associations with an FDR < 5% are reported.

### Gene-based association analyses

Gene-based association analyses were carried out using S-MultiXcan [26] implemented in the CTG-VL [31], which conducts a test of association between phenotypes and gene expression levels predicted by data derived from the GTEx project [56]. In this study, we performed S-MultiXcan using prediction models of all the brain tissues available from the GTEx project. This included amygdala, anterior cingulate cortex BA24, caudate basal ganglia, cerebellar hemisphere, cerebellum, brain cortex, frontal cortex BA9, hippocampus, hypothalamus, nucleus accumbens basal ganglia, putamen basal ganglia, spinal cord cervical c-1, substantia nigra. As a total of 14,501 genes expressed in different brain tissues were tested, the Bonferroni-corrected significance threshold was set at P = 3.44 × 10^−6^.

### Genetic correlations

In order to test whether handedness shares a genetic background with other complex traits with GWAS summary data available, we used CTG-VL [31] which implements LD score regression and contains a large database of summary GWAS statistics. In total, we assessed the genetic correlation of left-handedness and ambidexterity with 1,349 different traits. After accounting for multiple testing, our Bonferroni corrected significance threshold was P < 3.7 × 10^−5^.

### Heritability estimates

In order to estimate the proportion of phenotypic variance explained by SNPs we used two statistical methods. Restricted Maximum Likelihood (REML), implemented in BOLT-LMM was used to estimate the variance explained by additive effects of genotyped SNPs (h^2^_g_) [57]. Using a prevalence estimate of 10% the observed h^2^_g_ was transformed to a SNP heritability on an unobserved continuous liability scale [58]. LD-score regression was used to estimate the variance explained by all the SNPs using the GWAS summary statistics. Similarly to REML, the observed h^2^_g_ was transformed to the liability scale using a prevalence estimate of 10%.

In order to estimate the narrow-sense heritability we fit a variance components model to estimate the proportion of phenotypic variance attributable to additive genetic effects (A), shared environmental effects (C), and environmental effects (E) [28]. We modelled the genetic sharing between close relative pairs using identity by descent (IBD) information on 20,277 sibling pairs (0.65>IBD>0.35) and 49,788 relative pairs with 0.3>IBD>0.8 to estimate trait heritability. In the model, we also estimated a variance component due to a shared environment (siblings only) that made siblings potentially more similar in terms of handedness, and a unique environmental component, that did not contribute to similarity between relative pairs. Variance components were estimated using Maximum Likelihood using the OpenMx package [59].

## Acknowledgements

This research was conducted using the UK Biobank Resource (application number 12703). Access to the UK Biobank study data was funded by the University of Queensland (Early Career Researcher Grant 2014002959 to N.M.W). DME is funded by a National Health and Medical Research Council Senior Research Fellowship (APP1137714). GCP is funded by an Australia Research Council Discovery Early Career Researcher Award (DE180100976). CML is supported by the Li Ka Shing Foundation, WT-SSI/John Fell funds and by the NIHR Biomedical Research Centre, Oxford, by Widenlife and NIH (5P50HD028138-27). MMcC funding support for this work comes from Wellcome (090532, 106130, 098381, 203141, 212259, 095101, 200837, 099673/Z/12/Z) and NIHR (NF-SI-0617-10090). NMW is supported by an Australian National Health and Medical Research Council Early Career Fellowship (APP1104818). SEM was funded by an NHMRC Senior Reseach Fellowship (APP1103623). IB is funded by Wellcome (WT206194).

We thank the research participants of 23andMe for making this study possible. Members of the 23andMe Research Team are: Michelle Agee, Adam Auton, Robert K. Bell, Katarzyna Bryc, Sarah L. Elson, Pierre Fontanillas, Nicholas A. Furlotte, Barry Hicks, Karen E. Huber, Ethan M. Jewett, Yunxuan Jiang, Aaron Kleinman, Keng-Han Lin, Nadia K. Litterman, Matthew H. McIntyre, Kimberly F. McManus, Joanna L. Mountain, Elizabeth S. Noblin, Carrie A.M. Northover, Steven J. Pitts, G. David Poznik, J. Fah Sathirapongsasuti, Janie F. Shelton, Suyash Shringarpure, Chao Tian, Vladimir Vacic, Xin Wang, Catherine H. Wilson.

We are extremely grateful to all the families who took part in the ALSPAC study, the midwives for their help in recruiting them, and the whole ALSPAC team, which includes interviewers, computer and laboratory technicians, clerical workers, research scientists, volunteers, managers, receptionists and nurses. The UK Medical Research Council and Wellcome Trust (Grant Ref: 102215/2/13/2) and the University of Bristol provide core support for ALSPAC. GWAS data was generated by Sample Logistics and Genotyping Facilities at Wellcome Sanger Institute and LabCorp (Laboratory Corporation of America) using support from 23andMe.

The Netherlands Twin Register acknowledges funding from the Netherlands Organization for Scientific research (NWO), including NWO-Grant 480-15-001/674: Netherlands Twin Registry Repository and the Biobanking and Biomolecular Resources Research Infrastructure (BBMRI –NL, 184.021.007 and 184.033.111); KNAW Academy Professor Award (PAH/6635 to DIB); Amsterdam Public Health (APH) and Neuroscience Campus Amsterdam (NCA); European Community 7th Framework Program (FP7/2007-2013): ENGAGE (HEALTH-F4-2007-201413) and ACTION (9602768). The Rutgers University Cell and DNA Repository cooperative agreement (NIMH U24 MH068457-06); Collaborative study of the genetics of DZ twinning (NIH R01D0042157-01A1); Developmental Study of Attention Problems in Young Twins (NIMH, RO1 MH58799-03); Major depression: stage 1 genome-wide association in population-based samples (MH081802); Determinants of Adolescent Exercise Behavior (NIDDK R01 DK092127-04); Grand Opportunity grants Integration of genomics and transcriptomics (NIMH 1RC2MH089951-01) and Developmental trajectories of psychopathology (NIMH 1RC2 MH089995); and the Avera Institute for Human Genetics, Sioux Falls, South Dakota (USA).

The generation and management of GWAS genotype data for the Rotterdam Study (RS I, RS II, RS III) was executed by the Human Genotyping Facility of the Genetic Laboratory of the Department of Internal Medicine, Erasmus MC, Rotterdam, The Netherlands. The GWAS datasets are supported by the Netherlands Organisation of Scientific Research NWO Investments (nr. 175.010.2005.011, 911-03-012), the Genetic Laboratory of the Department
of Internal Medicine, Erasmus MC, the Research Institute for Diseases in the Elderly (014-93-015; RIDE2), the Netherlands Genomics Initiative (NGI)/Netherlands Organisation for Scientific Research (NWO) Netherlands Consortium for Healthy Aging (NCHA), project nr. 050-060-810. We thank Pascal Arp, Mila Jhamai, Marijn Verkerk, Lizbeth Herrera and Marjolein Peters, MSc, and Carolina Medina-Gomez, MSc, for their help in creating the GWAS database, and Karol Estrada, PhD, Yurii Aulchenko, PhD, and Carolina Medina-Gomez, PhD, for the creation and analysis of imputed data. We would like to thank Dr. Karol Estrada, Dr. Fernando Rivadeneira, Dr. Tobias A. Knoch, Marijn Verkerk, Anis Abuseiris, The Rotterdam Study is
funded by Erasmus Medical Center and Erasmus University, Rotterdam, Netherlands Organization for the Health Research and Development (ZonMw), the Research Institute for Diseases in the Elderly (RIDE), the Ministry of Education, Culture and Science, the Ministry for Health, Welfare and Sports, the European Commission (DG XII), and the Municipality of Rotterdam. The authors are very grateful to the study participants, the staff from the Rotterdam Study (particularly L. Buist and J.H. van den Boogert) and the participating general practitioners and pharmacists.

We thank all study participants as well as everybody involved in the Helsinki Birth Cohort Study (HBCS). Helsinki Birth Cohort Study has been supported by grants from the Academy of Finland, the Finnish Diabetes Research Society, Folkhälsan Research Foundation, Novo Nordisk Foundation, Finska Läkaresällskapet, Juho Vainio Foundation, Signe and Ane Gyllenberg Foundation, University of Helsinki, Ministry of Education, Ahokas Foundation, Emil Aaltonen Foundation.

In addition, we thank the participants and staff of the Nurses’ Health Study and the Health Professionals Follow-up Study, for their valuable contributions and Channing Division of Network Medicine, Department of Medicine, Brigham and Women’s Hospital, Harvard Medical School, Boston, MA. This work was supported in part by NIH R01 CA49449, P01 CA87969, UM1 CA186107, and UM1 CA167552.

The KORA study was initiated and financed by the Helmholtz Zentrum München-German Research Center for Environmental Health, which is funded by the German Federal Ministry of Education and Research (BMBF) and by the State of Bavaria. Furthermore, KORA research was supported within the Munich Center of Health Sciences (MC-Health), Ludwig-Maximilians-Universität, as part of LMUinnovativ.

We also thank funding for the ENGAGE (European Network for Genetic and Genomic Epidemiology) Consortium by the European Commission (FP7-HEALTH-F4-2007 (201413)).

Finally, we thank the participants of all the cohorts for their valuable contribution.

This publication is the work of the authors and DME will serve as guarantor for the contents of this paper.

## Conflict of interest statement

NE, DH, JT, and members of the 23andMe Research Team are employees of 23andMe, Inc., and hold stock or stock options in 23andMe. SHM, KS, HS, SS and GT are employees of deCODE Genetics/Amgen.

MMcC is a Wellcome Senior Investigator and an NIHR Senior Investigator. The views expressed in this article are those of the author(s) and not necessarily those of the NHS, the NIHR, or the Department of Health. He has served on advisory panels for Pfizer, NovoNordisk, Zoe Global; has received honoraria from Merck, Pfizer, NovoNordisk and Eli Lilly; has stock options in Zoe Global; has received research funding from Abbvie, Astra Zeneca, Boehringer Ingelheim, Eli Lilly, Janssen, Merck, NovoNordisk, Pfizer, Roche, Sanofi Aventis, Servier, Takeda. As of June 2019, he is an employee of Genentech, and holder of Roche stock.

All other authors report no conflicts of interest.

The 23andMe summary statistics used in this study will be made available to qualified researchers who enter into a data transfer agreement with 23andMe that protects participant privacy. Interested investigators should visit research.23andme.com/dataset-access to submit a request.

